# History-dependent perturbation response in limb muscle

**DOI:** 10.1101/509646

**Authors:** Thomas Libby, Chidinma Chukwueke, Simon Sponberg

## Abstract

1

Muscle mediates movement but movement is typically unsteady and perturbed. Muscle is known to behave non-linearly and with history dependent properties during steady locomotion, but the importance of history dependence in mediating muscles function during perturbations remains less clear. To explore muscle’s capacity to mitigate perturbations, we constructed a series of perturbations that varied only in kinematic history, keeping instantaneous position, velocity and time from stimulation constant. We discovered that muscle’s perturbation response is profoundly history dependent, varying by four fold as baseline frequency changes, and dissipating energy equivalent to ~ 6 times the kinetic energy of all the limbs (nearly 2400 W *Kg*^−1^). Muscle’s energy dissipation during a perturbation is predicted primarily by the force at the onset of the perturbation. This relationship holds across different frequencies and timings of stimulation. This history dependence behaves like a viscoelastic memory producing perturbation responses that vary with the frequency of the underlying movement.

**Summary Statement:** The response of muscles to rapid, identical strain perturbations is history dependent, but is captured by a viscoelastic model with memory. Muscle function during perturbations therefore depends on locomotor frequency.

## 2 Introduction

Muscle produces, dissipates, stores, returns, and transits mechanical energy to adopt diverse functions during locomotion (Dickinson et al., 2000). Even the same muscle can adopt different functions in unsteady or perturbed conditions (Biewener and Daley, 2007; Azizi and Roberts, 2010). A single muscle in the leg of cockroach normally dissipates energy during steady-state running (Ahn et al., 2006; Full et al., 1998). Yet when the animal is perturbed, neural feedback can categorically switch the muscle’s function from one stride to the next (Sponberg, Spence, Mullens and Full, 2011). Under unsteady conditions the muscle can dissipate more than ten times the energy that it does in steady state or convert its function to that of non-linear motor (Sponberg, Libby, Mullens and Full, 2011). It remains challenging to predict function from the quasi-static length-tension and force-velocity relationships, especially under unsteady conditions. Nonetheless such conditions likely pose greater performance demands than steady-state.

Strain history-dependent muscle properties are well known to affect muscle’s stress development. These properties include force depression during shortening and force enhancement during lengthening. While the specific mechanisms for history dependence remain controversial and are likely multifaceted (Rassier, 2012), there are established consequences for steady, transition, and impulsive behaviors (Josephson, 1999; Roberts and Azizi, 2011; Herzog et al., 2015; Nishikawa, 2016). However, muscle function during perturbations during movement is much less explored. Perturbations around steady-state, typically tetanic conditions are ubiquitous and simple material models like a viscoelastic Voigt body or a three component Hill model can typically capture muscle behavior in these cases (*e.g*. (Kirsch et al., 1994; Zajac, 1989; Cannon and Zahalak, 1982). However, dynamic conditions can create unexpected shifts in muscle performance (Robertson and Sawicki, 2015). During running, muscle experiences large rapid perturbations against a background strain trajectory where history has the potential to alter function. This modulation could have profound effects on muscle’s response to unsteady perturbations encountered during running, including slips or impacts with the substrate. Does history significantly modulate work output during rapid perturbations to periodic trajectories and can we reconcile any non-linearity with the simple material models that capture perturbations in static conditions?

To test these ideas we construct a systematic perturbation to a cockroach limb muscle that can reveal the importance of history on transient behavior and identify simple predictors of function (Fig. 1a). To do this we maintain the same Hill-model contractile properties (stimulation, strain trajectory, and velocity), while changing the strain history leading up to a perturbation. We modify this history by changing the frequency of background strain. We hypothesize that history dependence modulates muscle’s mechanical response to rapid perturbations, but that the response will be predictable from the components of an active viscoelastic system. If history dependence has a functionally relevant consequence for rapid mechanical perturbations then muscle work during the perturbation should vary systematically with history. If this history dependence matters for locomotion then the modulation produced should be significant in light of the mechanical power required to alter limb movement.

**Figure 1:**
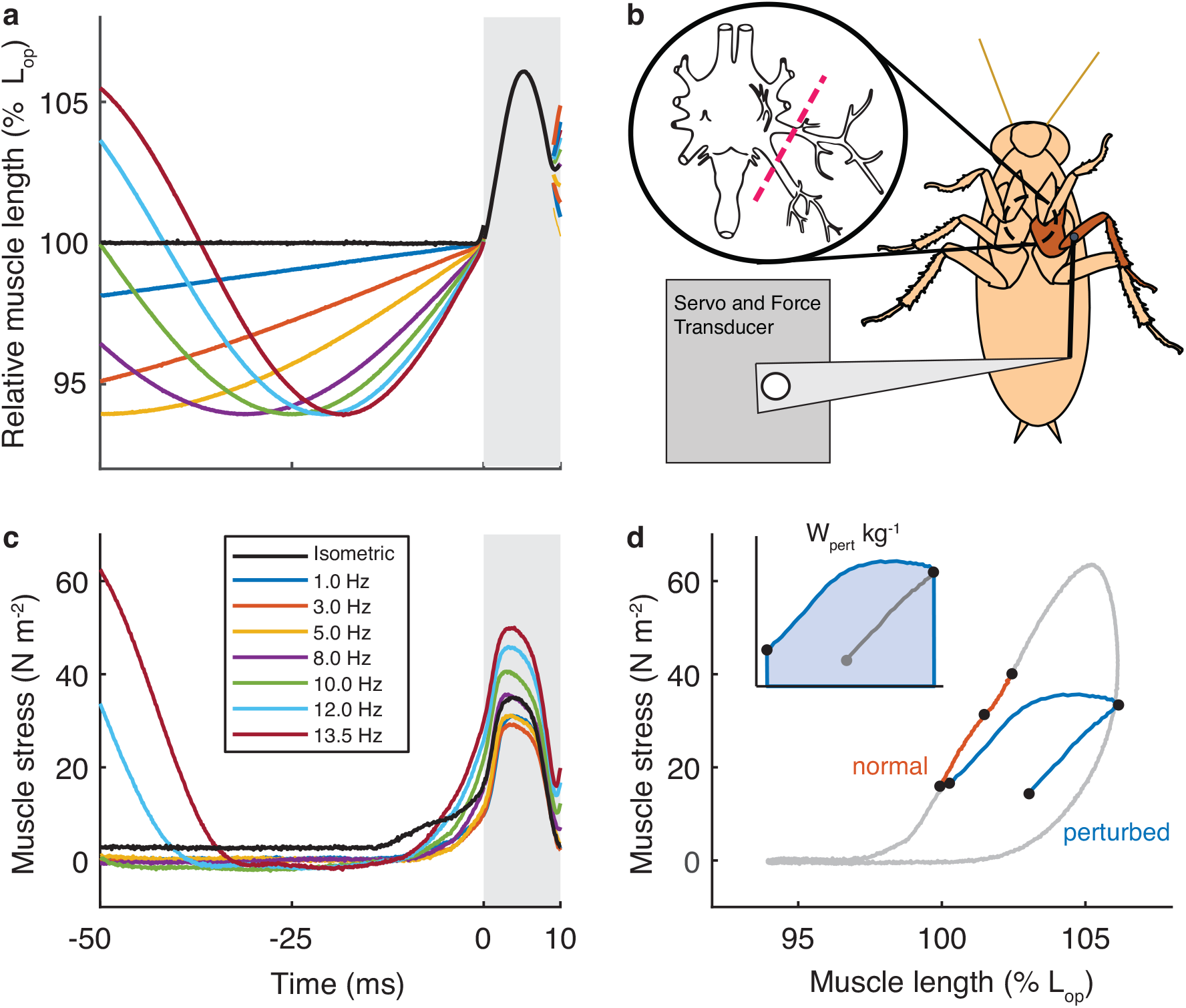
Identical eccentric perturbations (grey region) were applied during sinusoidal strain cycles of 1 Hz to 13.5 Hz (a). The intact joint workloop preparation (b) extracellularly stimulates muscles in the deinnervated limb (inset shows ganglion nerves cut adapted from (Pipa and Cook, 1959)). After subtracting out the passive joint work throughout the cycle (Sponberg, Libby, Mullens and Full, 2011), muscle stress (c) was integrated over strain for the eccentric portion of the perturbation to calculate work, *W_pert_* (d).

## 3 Materials and methods

Cockroaches (*Blaberus discoidalis*) were housed on a 12:12 light-dark cycle and fed dog chow *ad libitum*. We targeted the ventral femoral extensor of the middle leg (muscle 137), which is not the extensor that primarily powers limb extension, but rather a control muscle implicated in perturbation responses to locomotion on rough terrain (Full et al., 1998; Ahn and Full, 2002; Sponberg and Full, 2008). Instead of isolated muscle work loops (Josephson, 1985), we used intact joint work loops following previous methods (Fig. 1b, (Sponberg, Libby, Mullens and Full, 2011)). In brief, all motor neurons innervating the middle leg are severed at the mesothoracic ganglion by surgical ablation of Nerves 3, 4, 5, and 6. The limb is then mounted on a custom restraint stage and the coxa immobilized with epoxy. A muscle ergometer (Aurora Scientific 305C) attaches to the femur near the coxa femur joint via a two pin joint that allows for rotation. The femur and more distal segments are removed and the target muscle is activated via implant bipolar silver wire electrodes. The moment arm, pivot point, and linear relationship between joint angle and muscle strain are taken from prior work on this muscle. During steady-state work loop and imposed perturbations, the ergometer prescribed joint trajectories and simultaneously measured force. While this precludes the muscle from dynamically interacting with a load perturbation, it enabled us to use comparable conditions that vary only in history.

Besides the advantage of preserving the animals nutrient and oxygen supply to the target muscle, the intact joint work loop allows us to estimate the total passive work done on the joint. Here, muscle work loops are the active component of the work, calculated by measuring a passive work loop under identical strain conditions (including the perturbation) and subtracting the force measured in the passive trial from that of the active trial. The remaining force signal can be converted to muscle force (through the lever arm ratios) or used to calculate muscle work. Prior work (Sponberg, Libby, Mullens and Full, 2011) validated this approach as reflective of the work output and muscle function reported in more traditional partially isolated, muscle work loops with direct neural stimulation (Full et al., 1998; Ahn et al., 2006).

Steady-state work loop conditions reflected strain trajectory from *in vivo* 10 Hz running conditions and included three muscle potentials (spikes) of stimulation at 10 ms interspike interval and 0.5 ms duration applied at the onset of shortening. Stimulus voltage for each preparation was tuned to the minimum voltage needed to elicit a plateaued twitch response plus one volt. Perturbations were imposed halfway through shortening (stance) phases. We constructed 10 ms (100 Hz) half-sine perturbations of amplitude equal to the stride amplitude. These strain perturbations were not summed with background periodic strain but rather pasted in place, so that kinematics were exactly identical across all history conditions (Fig. 1a). The initial and terminal 1 ms of the perturbation was smoothed into the underlying strain trajectory using a linear ramp filter to prevent discontinuities in velocity.

We modified kinematics and timing of stimulus to test the effect of history on perturbation response. Perturbation kinematics remained constant in all conditions. To preserve Hill determinants during the perturbation, we varied the phase of stimulation so that the spike train always began 20 ms before the onset of perturbation (hence the stimulation in these altered conditions did not reflect natural timing). We changed the phase of the background work loop so that perturbations occurred both mid-stance and mid-swing, and we varied cycle frequency from 1 to 13.5 Hz, which was the fastest condition where we could maintain accurate perturbation conditions. To test the generality of our results, we repeated these conditions at different stimulation timings varying from 10 to 40 ms.

## 4 Results and discussion

### 4.1 Muscle’s perturbation response is history-dependent

Muscles absorbed energy during all perturbations (Fig. 1c). We characterized the muscle’s responses by the mechanical energy dissipated during the stretch portion of the perturbation (the area under the force-length curve (Fig. 1d). Muscles absorbed 23.6 ± 4.6 *Jkg*^−1^ during perturbations applied against a 10 Hz pre-lengthened background. A Hill-type contractile unit would respond with identical force profiles to the perturbation regardless of kinematic history. By contrast, we found that active muscle stress and energy dissipation increased strongly as we increased frequency under pre-lengthened conditions, and decreased with frequency under pre-shortened conditions (Fig. 2a). Energy absorption varied almost 4-fold over the range of frequencies we tested. At the typical running stride frequency of 10 Hz, dissipation almost tripled if the muscle was perturbed in identical ways during shortening and lengthening.

**Figure 2:**
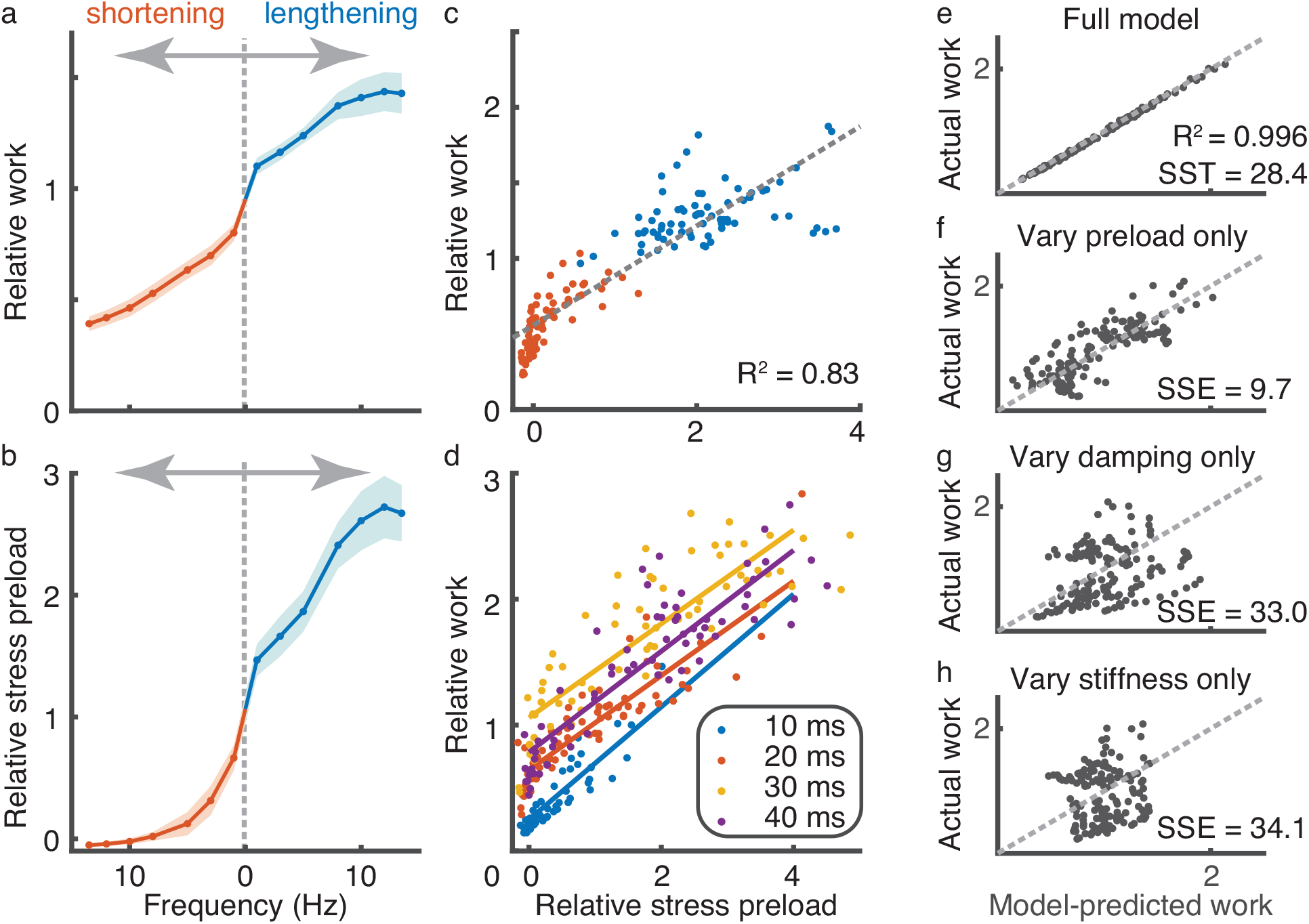
Work during perturbation (a) and preload stress (b) both vary significantly and monotonically with frequency, resulting in a correlated response (c). This relationship persists if stimulation varies by changing how long before the perturbation the muscle is electrically stimulated (d). To normalize across animals, work is taken relative to the average of the two 1 Hz perturbation conditions which most closely approximate the isometric work response. Relative stress is scaled to the isometric response to *in vivo* stimulation conditions (3 spikes, 10 ms interspike interval). Preload refers to stress developed immediately prior to the onset of the perturbation. A 3-parameter viscoelastic model with memory (e) predicts muscle’s work during the perturbation (Fig. 1d). While all three parameters contribute to the full model, preload (f) has a predictive effect on its own, unlike damping (g) and stiffness (h). A sum of square errors (SSE) greater than the total sum of squares (SST) indicates a non-predictive model (g & h).

### 4.2 Force at onset of perturbation determines work

If preloading is the primary determinant of the muscle’s variable perturbation response, then the force at the onset of the perturbation should parallel the change in dissipative work. Indeed, muscle stress prior to perturbation onset showed a pattern of frequency dependence similar to work (Fig. 2b) and the two are highly correlated (Fig. 2c, *R*^2^ = 0.83, *p* < 10^−6^). Despite the sharp deviations from an otherwise monotonic work relationship on pre-stretch at 15 Hz, the stress at onset of these perturbations still recapitulated the changes in absorption. At the fastest shortening velocities (left side of Fig. 2 a,c), work continues to fall off even after preload stress has reached zero likely because of the need to take up slack from the transition of rapid shortening to lengthening.

To further test the robustness of this relationship, we repeated the frequency and phase experiments while varying the timing of muscle stimulation prior to perturbation. Muscle stress prior to perturbation near static conditions increased almost 10 fold as timing advanced from 10 *ms* to 40 *ms*, but both the frequency dependency and the correlation between stress and work persisted across stimulation conditions (Fig. 2d, *R* > 0.85, *p* < 10^−6^ for all timings).

A Hill-type contractile element model fails to predict muscle’s perturbation response, even when time scales are quite rapid. Instead we support the hypothesis that history dependence tunes muscle’s mechanical response to perturbations. Muscle’s response to rapid stretches is known to have viscoelastic properties (Kirsch et al., 1994; Zajac, 1989); here we show that the context in which a perturbation occurs, meaning the muscle’s force history, modulates these properties and shapes muscle’s function even on short time scales. While Hill-like contractile elements fail to directly predict the functional modulation during the perturbation, they do play a role because the pre-perturbation forces follows a classic force-velocity curve (Fig. 2b), velocity is proportional to frequency). Despite history varying muscle’s mechanical work (Fig. 2a), the behavior of muscle is nonetheless predictable. This relationship holds regardless of whether the muscle is pre-shortened or pre-lengthened and across a range of frequencies and stimulation spanning natural running (Fig. 2c, d).

### 4.3 A viscoelastic model with memory captures muscle’s rapid perturbation response

Since muscle force-length behavior during the perturbations appeared viscoelastic (Fig. 1d), we fit a parallel spring-damper (Voigt) model with a variable preload offset to the perturbation data. Changes in dissipative work against history could arise from I) a change in stiffness, II) increased viscous damping, or III) a change in the preload force. The full model fit these parameters separately for each condition (phase and frequency) and strongly predicted observed energy absorption (Fig. 2e, *R*^2^ > 0.99). To examine which parameters were most predictive, we tested three models which each allowed only one parameter to vary (with the other two set to the average across all frequencies/phases). Doing so reduced the variance accounted for in all cases, but only when preload varied did the model retain any predictive ability (Fig. 2f-h).

The model property that best explained the functional variation was preload, rather than stiffness or damping. In prior studies with small sinusoid perturbation, muscle stiffness varied with frequency (Cannon and Zahalak, 1982; Kirsch et al., 1994). Here, variable stiffness does not account for the major differences in perturbation work, likely because total strain, velocity and prior stimulation were kept the same. During perturbations to steady state conditions, work also varies with pre-perturbation force (Kirsch et al., 1994). We show that this adjustment, equivalent to changing the set point of a viscoelastic model, is the critical feature for dynamic conditions and can account for differences in perturbations responses relevant to locomotion. Our results are consistent with a viscoelastic memory effect, which typically arises from stored energy in elastic structures that cannot relax instantaneously.

What are the potential mechanisms for this non-linearity? Series elastic elements (SSEs) can modulate the state of the muscle fibers, contributing to energy storage and return, power amplification, or dissipation (Roberts and Azizi, 2011). Yet SSE’s role in rapid perturbations is less explored. Our results might be explained by a contractile element with a series elastic component. However, the viscoelastic properties of insect apodeme and the exoskeletal attachment are typically an order of magnitude stiffer than vertebrate tendon (Bennet-Clark, 1975; Zajac, 1989). They are also short and would require ~ 60% strain to account for the perturbation if the muscle remained isometric. It is likely that the muscle must be significantly involved in modulating the preloading and hence the work done during the perturbation. Other contributors to series elasticity could also play a role.

Nonuniformity in sarcomere strain and force production (Rassier and Herzog, 2004) is a likely contributor to history dependence, but it is unlikely to be the only explanation because history dependence typically manifests in single sarcomeres (Leonard et al., 2010; Rassier, 2012). There is also growing appreciation that components of the muscle lattice other than actin and myosin might contribute to history dependent phenomena (Rassier, 2012; Herzog et al., 2015; Nishikawa, 2016). Titin and other large structural proteins have been implicated in history dependent properties in muscle. Calcium-dependent binding of the N2A domain effectively changes the stiffness and offset of the spring-like PEVK domain (Herzog et al., 2015). Titin also is suggested to have a further role in force generation via active winding of titin around the thin filaments (Nishikawa, 2016; Lindstedt and Nishikawa, 2017). Other components might play a role as well, especially regulatory elements that are strain-dependent like tropomyosin (Tanner et al., 2012; Holt and Williams, 2018).

### 4.4 Muscle’s mechanical behavior during perturbations is significant for locomotion

Regardless of mechanism, the history-dependent modulation of work during a perturbation would only have meaningful consequences for locomotion if the overall change is significant in the context of muscle, joint, limb, and body. Surprisingly this capacity is substantial at all scales even in a relatively small muscle (Table 1). The energy absorbed by the muscle during the perturbation we applied is at least comparable to the kinetic energy of all the limbs (Kram et al., 1997) and at most could absorb the center of mass kinetic energy of a 3 *g* animal running at 20 *cm/s*! Despite being about 1/10^*th*^ the mass of the animal’s largest femoral extensor, muscle 137 absorbed more energy during a single perturbation at 10 Hz than the larger muscle produces during running (Ahn and Full, 2002). Muscle 137 absorbed about threefold more energy than the entire joint did during the same perturbation.

**Table 1:** Selected statistics (means ± 95% CI) and comparison measures from earlier work. Reference stress/work was used to normalize stress/work across conditions in Fig. 2. Reference work was the average across pre-lengthened-shortened conditions at 1 Hz.

|  |  |
| --- | --- |
| Average reference stress (isometric, 20 ms after first spike) | $17.3 \pm 5.8 \text{ Ncm}^{-2}$ |
| Average reference perturbation work | $-40.2 \pm 8.2 \mu J$ |
| Perturbation work at 10 Hz (eccentric history) | $-55.8 \pm 11.5 \mu J$ |
| Passive joint perturbation work | $-17.6 \pm 7.1 \mu J$ |
| Average specific dissipation power at 10 Hz | $-2360 \pm 798 \text{ Wkg}^{-1}$ |
| Muscle 137 *in situ* negative work Full et al. (1998) | $-9 \mu J$ |
| Muscle 137 *in situ* max positive work, 6 spikes Sponberg, Libby, Mullens and Full (2011) | $25 \mu J$ |
| Muscle 137 *in situ* max negative work, 8 spikes Sponberg, Libby, Mullens and Full (2011) | $-45 \mu J$ |
| Muscle 177c *in situ* positive work Ahn and Full (2002) | $74 \mu J$ |
| Peak kinetic energy of all limbs, 20 cm/s running Kram et al. (1997) | $8 \mu J$ |

This single cockroach muscle therefore serves as an example of the versatile control role a muscle can adopt. At steady state during running, it typically dissipates a small amount of energy during the swing period of each stride (Full et al., 1998). Its steady-state work is far below its capacity to either dissipate energy or do mechanical work when neural feedback modulates its function during locomotion (Sponberg, Spence, Mullens and Full, 2011). Neural feedback can also turn this muscle into a motor assisting in obstacle traversal or turning (Sponberg, Spence, Mullens and Full, 2011). Our results here indicate another control function: open-loop tuning of the limb’s response to disturbances. Cockroaches locomote with stride frequencies that vary over a wide band (at least 1 – 12*Hz*); therefore the time available to stabilize perturbations decreases dramatically as running speed increases (Sponberg and Full, 2008). Even with constant timing of stimulation, muscle 137’s dissipative capacity during leg swing increases almost 50% over this frequency range. Unlike impulsive behaviors that can do work over a relatively long time (Ilton et al., 2018), both muscle’s dissipative power and total work matter for an effective response during perturbations.

But with the capacity to dissipate 2400 *W/kg*, a muscle need not be large to absorb energy quickly. Rapid movements in animals at many scales challenge sensorimotor bandwidth (More and Donelan, 2018). Context-dependent muscle behavior during perturbations could play an important role in stabilizing high-speed movements, especially if the animal can use anticipatory or feedforward control to tune muscle’s perturbation response.

Natural perturbations are unlikely to be prescribed kinematic deviations, although rigid obstacles such as on rough terrain could produce these. Exploring unsteady muscle function with perturbed work loops coupled to impulsive forces or simulated loads could lead to a more complete picture of perturbation responsiveness is specific muscles (Robertson and Sawicki, 2015). However, our approach of prescribed perturbations imposed at different frequencies isolated the effect of history dependence and showed that stretch enhancement and shortening depression have significant consequences for muscle’s function during perturbations.

## 5 Acknowledgements

The authors would like to thank Tom Daniel and C. Dave Williams for helpful comments regarding this manuscript.

## 8 Data accessibility

Data used in this study will be made available through the online repository Dryad upon publication.

## References

Ahn, A. N. and Full, R. J. (2002), ‘A motor and a brake: two leg extensor muscles acting at the same joint manage energy differently in a running insect’, Journal of Experimental Biology 205(3), 379–389.

Ahn, A. N., Meijer, K. and Full, R. J. (2006), ‘In situ muscle power differs without varying in vitro mechanical properties in two insect leg muscles innervated by the same motor neuron’, Journal of Experimental Biology 209(17), 3370–3382.

Azizi, E. and Roberts, T. J. (2010), ‘Muscle performance during frog jumping: influence of elasticity on muscle operating lengths’, Proceedings Of The Royal Society B-Biological Sciences 277(1687), 1523–1530.

Bennet-Clark, H. (1975), ‘The energetics of the jump of the locust Schistocerca gregaria.’, The Journal of experimental biology 63(1), 53–83.

Biewener, A. A. and Daley, M. A. (2007), ‘Unsteady locomotion: integrating muscle function with whole body dynamics and neuromuscular control.’, Journal of Experimental Biology 210(Pt 17), 2949–2960.

Cannon, S. C. and Zahalak, G. I. (1982), ‘The Mechanical Behavior of Active Human Skeletal-Muscle in Small Oscillations’, Journal of Biomechanics 15(2), 111–121.

Dickinson, M. H., Farley, C. T., Full, R. J., Koehl, M. A., Kram, R. and Lehman, S. (2000), ‘How animals move: an integrative view.’, Science 288(5463), 100–106.

Full, R. J., Stokes, D. R., Ahn, A. N. and Josephson, R. K. (1998), ‘Energy absorption during running by leg muscles in a cockroach’, Journal of Experimental Biology 201 (Pt 7)(7), 997–1012.

Herzog, W., Powers, K., Johnston, K. and Duvall, M. (2015), ‘A new paradigm for muscle contraction’, Frontiers in Physiology 6, 174.

Holt, N. C. and Williams, C. D. (2018), ‘Can Strain Dependent Inhibition of Cross-Bridge Binding Explain Shifts in Optimum Muscle Length?’, Integrative and comparative biology 58(2), 174–185.

Ilton, M., Bhamla, M. S., Ma, X., Cox, S. M., Fitchett, L. L., Kim, Y., Koh, J.-s., Krishnamurthy, D., Kuo, C.-Y., Temel, F. Z. et al. (2018), ‘The principles of cascading power limits in small, fast biological and engineered systems’, Science 360(6387).

Josephson, R. K. (1985), ‘Mechanical power output from striated-muscle during cyclic contraction’, Journal of Experimental Biology 114(1), 493–512.

Josephson, R. K. (1999), ‘Dissecting muscle power output’, Journal of Experimental Biology 202(23), 3369–3375.

Kirsch, R. F., Boskov, D. and Rymer, W. Z. (1994), ‘Muscle Stiffness During Transient and Continuous Movements of Cat Muscle: Perturbation Characteristics and Physiological Relevance’, IEEE Transactions on Biomedical Engineering 41(8), 758–770.

Kram, R., Wong, B. and Full, R. (1997), ‘Three-dimensional kinematics and limb kinetic energy of running cockroaches.’, Journal of Experimental Biology 200(13), 1919–1929.

Leonard, T. R., DuVall, M. and Herzog, W. (2010), ‘Force enhancement following stretch in a single sarcomere’, AJP: Cell Physiology 299(6), C1398–C1401.

Lindstedt, S. and Nishikawa, K. (2017), ‘Huxleys’ Missing Filament: Form and Function of Titin in Vertebrate Striated Muscle’, Annual Review of Physiology 79(1), 145–166.

More, H. L. and Donelan, J. M. (2018), ‘Scaling of sensorimotor delays in terrestrial mammals’, Proceedings of the Royal Society B: Biological Sciences 285(1885), 20180613.

Nishikawa, K. (2016), ‘Eccentric contraction: unraveling mechanisms of force enhancement and energy conservation’, Journal of Experimental Biology 219(2), 189–196.

Pipa, R. L. and Cook, E. F. (1959), ‘Studies on the hexapod nervous system. I. The peripheral distribution of the thoracic nerves of the adult cockroach, Periplaneta americana’, Annals of the Entomological Society of America 52, 695–710.

Rassier, D. E. (2012), ‘The mechanisms of the residual force enhancement after stretch of skeletal muscle: Non-uniformity in half-sarcomeres and stiffness of titin’, Proceedings of the Royal Society B: Biological Sciences 279(1739), 2705–2713.

Rassier, D. E. and Herzog, W. (2004), ‘Considerations on the history dependence of muscle contraction’, Journal Of Applied Physiology 96(2), 419–427.

Roberts, T. J. and Azizi, E. (2011), ‘Flexible mechanisms: the diverse roles of biological springs in vertebrate movement’, Journal of Experimental Biology 214(3), 353–361.

Robertson, B. D. and Sawicki, G. S. (2015), ‘Unconstrained muscle-tendon workloops indicate resonance tuning as a mechanism for elastic limb behavior during terrestrial locomotion.’, Proceedings of the National Academy of Sciences 112(43), E5891–8.

Sponberg, S. and Full, R. J. (2008), ‘Neuromechanical response of musculo-skeletal structures in cockroaches during rapid running on rough terrain.’, Journal of Experimental Biology 211(Pt 3), 433–446.

Sponberg, S., Libby, T., Mullens, C. H. and Full, R. J. (2011), ‘Shifts in a single muscle’s control potential of body dynamics are determined by mechanical feedback.’, Philosophical Transactions Of The Royal Society Of London Series B-Biological Sciences 366(1570), 1606–1620.

Sponberg, S., Spence, A. J., Mullens, C. H. and Full, R. J. (2011), ‘A single muscle’s multifunctional control potential of body dynamics for postural control and running.’, Philosophical Transactions Of The Royal Society Of London Series B-Biological Sciences 366(1570), 1592–1605.

Tanner, B. C. W., Daniel, T. L. and Regnier, M. (2012), ‘Filament Compliance Influences Cooperative Activation of Thin Filaments and the Dynamics of Force Production in Skeletal Muscle’, PLoS Computational Biology 8(5).

Zajac, F. E. (1989), ‘Muscle and tendon: properties, models, scaling, and application to biomechanics and motor control’, Crit Rev Biomed Eng 17(4), 359–411.

